# A comprehensive *in silico* investigation into the nsSNPs of *Drd2* gene predicts significant functional consequences in dopamine signaling and pharmacotherapy

**DOI:** 10.1101/2021.06.11.448090

**Authors:** Samia Sultana Lira, Ishtiaque Ahammad

## Abstract

DRD2 is a neuronal cell surface protein involved in brain development and function. Variations in the *Drd2* gene have clinical significance since DRD2 is a pharmacotherapeutic target for treating psychiatric disorders like ADHD and schizophrenia. Despite numerous studies on the disease association of single nucleotide polymorphisms (SNPs) in the intronic regions, investigation into the coding regions is surprisingly limited. In this study, we aimed at identifying potential functionally and pharmaco-therapeutically deleterious non-synonymous SNPs of *Drd2*. A wide array of bioinformatics tools was used to evaluate the impact of nsSNPs on protein structure and functionality. Out of 260 nsSNPs retrieved from the dbSNP database, initially 9 were predicted as deleterious by 15 tools. Upon further assessment of their domain association, conservation profile, homology models and inter-atomic interaction, the mutant F389V was considered as the most impactful. In-depth analysis of F389V through Molecular Docking and Dynamics Simulation revealed a decline in affinity for its native agonist dopamine and an increase in affinity for the antipsychotic drug risperidone. Remarkable alterations in binding interactions and stability of the protein-ligand complex in simulated physiological conditions were also noted. These findings will improve our understanding of the consequence of nsSNPs in disease-susceptibility and therapeutic efficacy.

## Introduction

Single Nucleotide Polymorphisms (SNPs) are the most common source of variance in the genome contributing greatly to phenotypic variation and several disease-association. In the case of the human genome, they are found in every 200–300 base pairs and represent 90% of genomic variation ^1–4^. SNPs in non-coding regions (for example, regulatory region, UTR, intron) can disrupt or modify various functions, such as interaction with miRNA and transcription factors, splicing, and ribosomal translation of mRNA ^5–9^. On the other hand, approximately 500,000 SNPs (on average 6 per gene) are located in the coding regions of the human genome ^4^. Among non-synonymous SNPs (nsSNPs), missense SNPs can cause the substitution of amino acids in protein sequences and thereby exert damaging or neutralizing effects. Alteration of sequences in the conserved region may result in deleterious effect by affecting protein function, structure, stability, translation, charge, hydrophobicity, geometry, dynamics, and inter/intra protein interactions ^10–12^. In fact, nsSNPs have been reported by numerous studies to influence the disease probability ^13–18^. Therefore, SNP association studies are useful for the characterization of phenotypes as well as insight into drug development and therapeutics of diseases concerned with the specific genome variation. However, owing to the large number of SNPs in *Drd2*, laboratory experimentation on the functional effects of these SNPs would be a costly and time-consuming laborious process. Therefore, computational screening of SNPs to shorten down the number of potential pathogenic ones is an essential step prior to experimental mutation analysis.

In this context, several *in silico* approaches have been largely utilized in the recent decade to predict the structural and functional influence of deleterious nsSNP in genes of interest. Several studies show that *in silico* algorithms have successfully predicted the significance of nsSNPs associated with specific genes ^19–26^. Though the accuracy of the tools can be questioned, they can still be used as a primary filter ^27^. Besides, combinatorial analysis of different algorithms makes the predicted effects of particular mutations more accurate. Moreover, refined analysis, such as molecular dynamics simulation enables precise evaluation of changes in protein structure, physicochemical properties, and interactions in a simulated environment ^28–32^.

DRD2 belongs to the G-protein-coupled receptor family and is a major mediator of the effects of dopamine. It is associated in the human brain with diverse functions such as reward mechanism, cognition, attention, movement, and neuroendocrine regulation, learning and memory. It has been reported to play a critical role in a number of clinical manifestations and neuro-psychiatric disorders such as addiction, neuroticism, parkinsonism, restless legs syndrome, schizophrenia, attention deficit hyper activity disorder (ADHD) and, bipolar disorder ^33–38^.

*Drd2* located at chromosome11q23.1 by alternative splicing leads to the generation of two isoforms ^39–41^. D2-long (D2L) is found to be expressed mostly post-synaptically, whereas D2-short(D2S) is a presynaptic isoform ^42^. Numerous genome-wide association studies (GWAS) have been carried out involving *Drd2* gene variants in intronic and regulatory regions, but not in the coding region ^43–50^. Although few studies on the coding sequence variant of *Drd2* are available ^51–60^, to our best knowledge, a comprehensive investigation is still absent.

With an aim to investigate the damaging effect of *Drd2* nsSNPs on disease predisposition and treatment responsiveness, in this study, we focused on the DRD2 protein structural and functional impairment upon mutation. For this purpose, a number of bioinformatics tools were employed to identify the most deleterious nsSNPs in *Drd2* and evaluate their effect on the gene product. After retrieval of the complete list of nsSNPs of *Drd2* from the dbSNP database, a number of prediction algorithms were put to use to detect pathogenic nsSNPs. Following analyses including conservation profile, domain position, post-translational modifications, we performed molecular docking and dynamics simulation for better understating of mutation impact in physiological condition.

## Results

The complete workflow employed in this study is summarized in Fig. 1.

**Figure 1.**
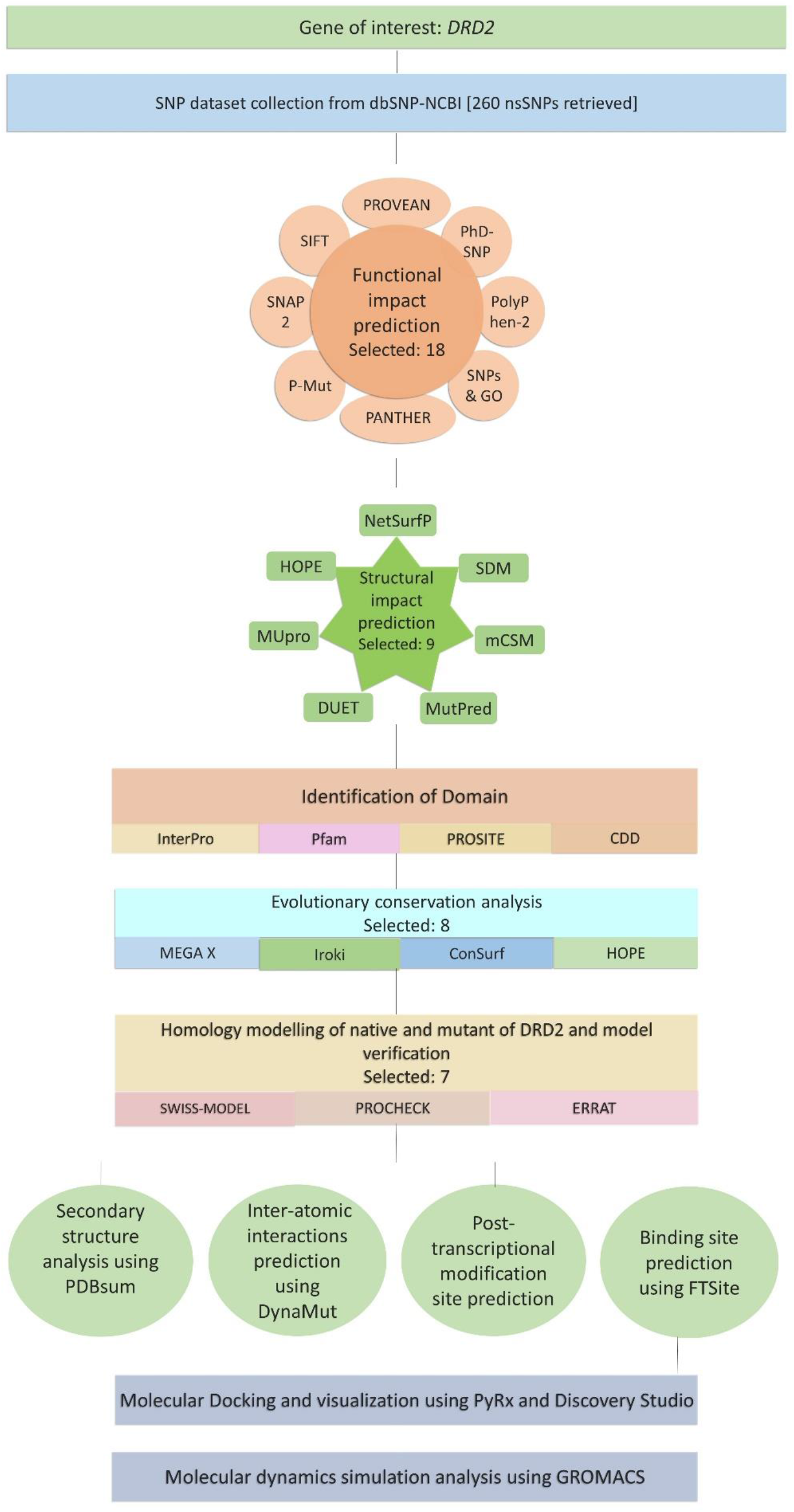
Schematic representation of the workflow of the study. The overall process can be summarized as a series of progressive filtration steps pursued for the identification of the most damaging nsSNPs of *Drd2* and subsequent in-depth analysis of one particular nsSNP.

### SNP data retrieval

The polymorphism information of *Drd2* was collected from the NCBI dbSNP database. Out of the total of 16,456 SNPs retrieved, the number of intronic, missense, synonymous, Non-coding variant and in-frame deletion amounted to 15,370 (93.40%), 238 (1.45%), 173 (1.05%), 139 (0.84%), and 2 (0.01%) respectively (Supplementary Table S1). Since the scope of the study is limited to nsSNPs of *Drd2*, only nsSNPs (total 260) were subjected to subsequent analysis.

### Functional impact prediction of *Drd2* nsSNPs

To determine the functional impact of nsSNPs on DRD2, a total of eight tools have been utilized (Supplementary Table S2). Out of the total of 260 nsSNPs, SIFT server predicted 119 to be functionally damaging, of which 27 had a tolerance index of 0. The PROVEAN server estimated 73 nsSNPs as deleterious out of all 260 nsSNPs submitted. The number of affecting ones anticipated by PolyPhen-2 is 139. Of the detrimental nsSNPs marked by PolyPhen-2, 102 were “probably damaging” (score 0.958-1.00) and 37 were “possibly damaging” (score 0.486-0.95). In addition, 215 nsSNPs were identified as damaging (171 of them are “probably damaging” and 44 are “possibly damaging”) by the server PANTHER-PSEP. The number of disease-associated nsSNPs predicted by SNAP-2, P-Mut, SNPs & GO and PhD-SNP are 145, 60, 50 and 70 respectively.

Among the nsSNPs subjected to analysis, 18 were unanimously predicted by 8 tools to have damaging effects and thereby were functionally significant. These 18 nsSNPs were taken into consideration for the next stage of filtering.

### Structural impact prediction of *Drd2* nsSNPs

Eighteen nsSNPs that had been predicted to be damaging by 8 different tools were then underwent structural impact analysis. For this purpose, seven servers were employed-MutPred, MUpro, NetSurfP, DUET, mCSM, SDM, and HOPE (Supplementary Table S3).

MutPred predicted all but one (Y37C) to be pathogenic while MUpro predicted that all the mutations for 2 (T119M, C443F) exerted a decreasing effect on the protein stability. NetSurfP predicts whether a residue is buried or exposed within the protein structure. It was observed that two of the mutants (A84T and R219C) shifted from exposed to buried and only one of the mutants (F389V) shifted from buried to exposed status when compared with the native structure. Furthermore, DUET predicted all but 4 and mCSM predicted all but 1 to be destabilizing whereas SDM predicted 11 of them to be destabilizing.

To investigate the effect of mutation on physico-chemical properties, hydrophobicity, intermolecular interaction as well as structural and functional changes, Project HOPE server was utilized. 17 mutations were predicted to cause a change in protein size and 9 mutations showed a change in charge. In terms of hydrophobicity, 8 showed modification while 2 completely lose their hydrophobicity. Moreover, 4 mutations were involved in cysteine bridges which were crucial for protein stability (C126W, R145C, R150C, and C399R) could impose a severe effect on the 3D structure of the protein.

Glycine residue at the 173^rd^ position of the native structure formed an unusual torsion angle, so mutant G173R was predicted to cause an incorrect conformation of the backbone. Proline on the native protein at 404^th^ position gave rise to special backbone conformation which might be disturbed by the mutation P404R. On the other hand, the native structure formed an H-bond with Gln368 and a salt bridge with Lys369. Mutation in the 368^th^ position, such as E368D, thereby, would hinder both the interactions. E358D, as well as R219C, could interrupt the interaction with Neurabin-2 which acted as a secondary messenger. F389V mutation being located within the agonist binding region, might disturb the functionality as a result of empty space in the core.

Finally, after the wide array of structural impact analyses by above-mentioned tools, 9 out of the 18 mutants were considered most impactful to the structure of the DRD2 protein and were selected for subsequent analyses. These mutations were C126W, R145C, R150C, G173R, R219C, E368D, F389V, C399R, and P404R.

### Identification of domain

To identify the domains in DRD2 and for mapping of nsSNPs, four different tools and databases-InterPro, Pfam, PROSITE, and CDD were used.

All 4 tools reported a seven-transmembrane domain in position 51-426, whereas CDD predicted the same domain to be located at the position 35-437. Our selected 9 nsSNPS were located within this domain.

### Conservation profile and evolutionary relationship analysis

Amino acid residues playing a critical role in numerous cellular processes such as genome stability tend to remain conserved despite evolutionary drift. Therefore, the intensity of residue conservation is often considered an indication of the importance of a position in maintaining protein stability and functionality ^61^. To inspect evolutionary conservation, phylogenetic analysis was carried out using the MEGA X server and was visualized by Iroki (Fig. 2) The result showed that the most closely related cousins of the DRD2 protein in *Homo sapiens* were their homologs in *Pan troglodytes* and *Pan paniscus*. Using the Bayesian method, the ConSurf web browser not only measured the conservation level of each residue in the protein but also revealed putative structural and functional residues. Out of 9 residues filtered out from upstream analysis, 4 structural(buried) and 3 functional(exposed) residues showed the highest and R145 showed a high degree of conservation (Fig.3, Supplementary Table S4). HOPE server predicted all except one of 9 residues to be very conserved. Mutation in P404 position was excluded from subsequent analysis since it was reported to have a low conservation profile by both ConSurf and HOPE servers.

**Figure 2.**
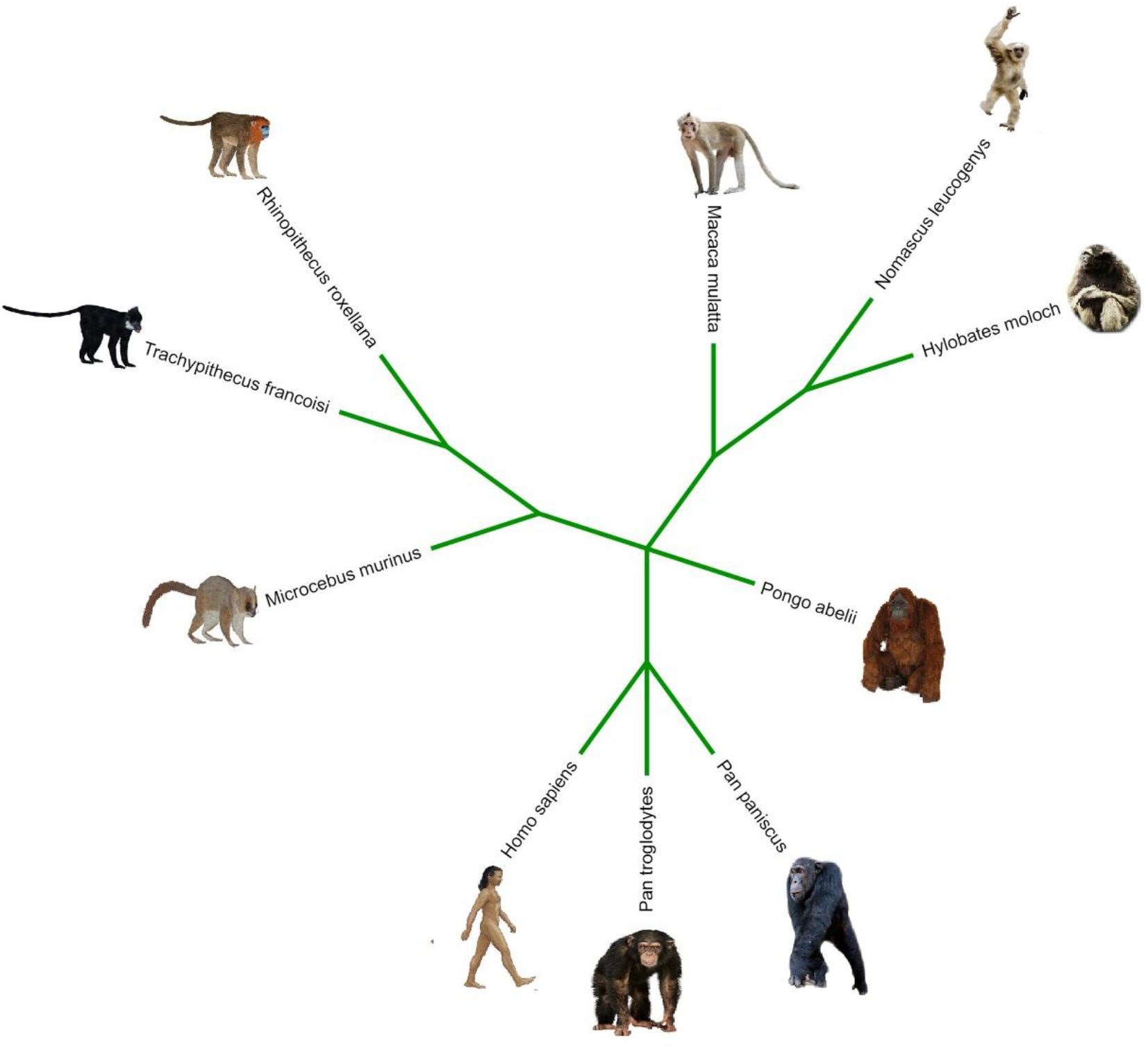
Graphical representation of evolutionary relationships between the human *Drd2* and its closest counterparts by Iroki server. The human DRD2 appears to be most closely related to its homologs in *Pan troglodytes* and *Pan paniscus*.

**Figure 3.**
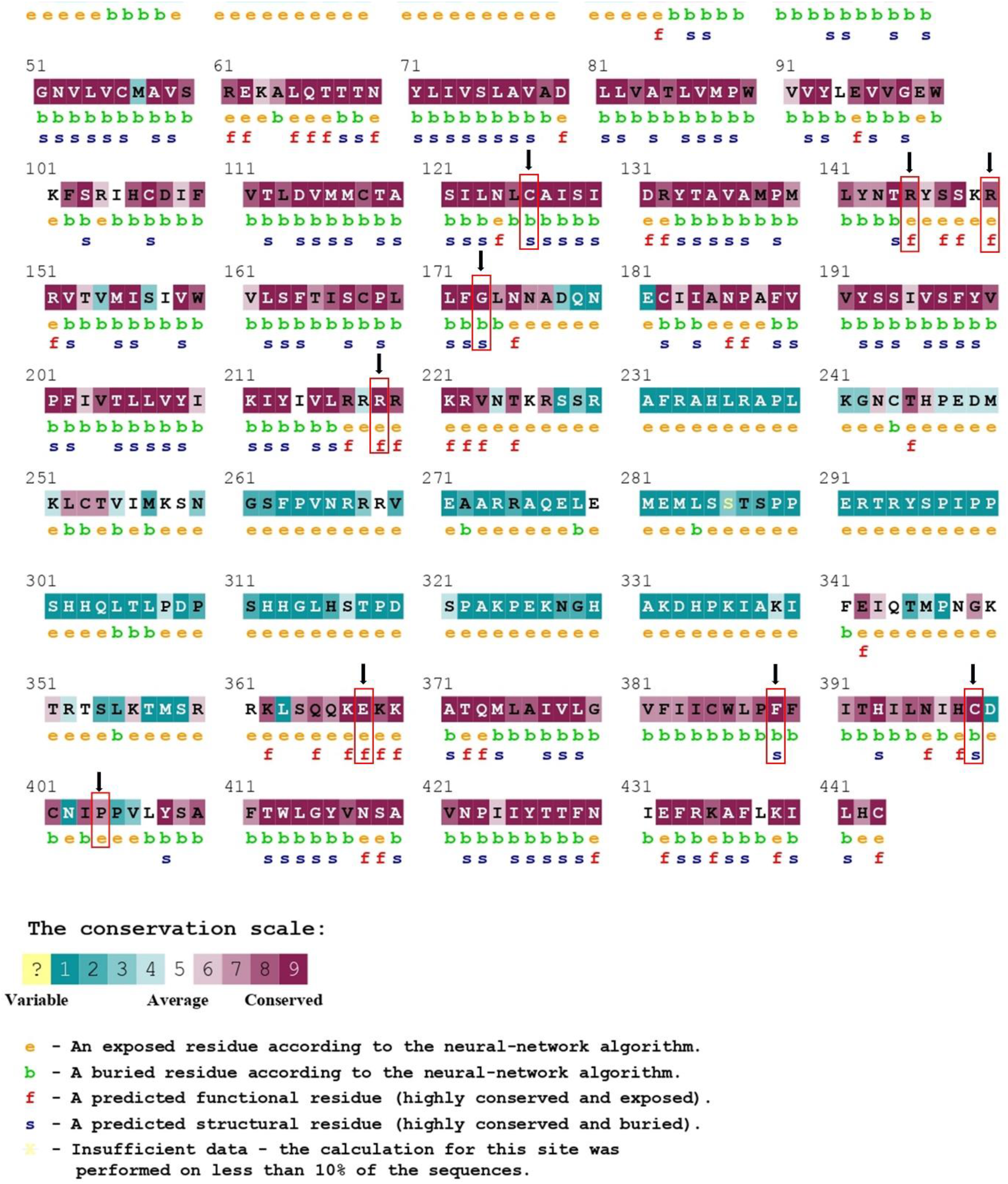
Evolutionary conservation profile of amino acid residues of DRD2 as predicted by ConSurf. Almost all the nsSNPs primarily evaluated as deleterious belonged to the highly conserved regions.

### Homology model verification and secondary structure analysis

In order to attain the 3D structure of native and 8 mutant proteins, SWISS-MODEL server was used. Model quality was verified by two different servers-PROCHECK and ERRAT (Supplementary Table S5). All the structures had up to the mark ERRAT quality factor. However, the Ramachandran plot generated by PROCHECK showed >90.0% residues in the most favored regions for all the structures except for G173R. (Supplementary Fig. S1) For sake of narrowing down the list, this mutation was excluded from further analysis.

Secondary structures of the native DRD2 and 7 mutants were analyzed and visualized using the tool PDBsum (Fig. 4) Except for E368D, all the mutants had a higher number of helices than the native one. In comparison to wild-type structure, five mutants (C126W, R145C, R150C, R219C, C399R) introduced one di-sulfide linkage in the same position while a subset of them (R145C and R219C) contained a shorter second di-sulfide bridge between the residues C399 and C401. Three mutants (C126W, R150C, and C399R) also included 3 A strands and 2 interspaced beta hairpins. Moreover, after the N259 position, F389V had more tightly packed β and γ turns while in the case of E368D, they were more interspersed.

**Figure 4.**
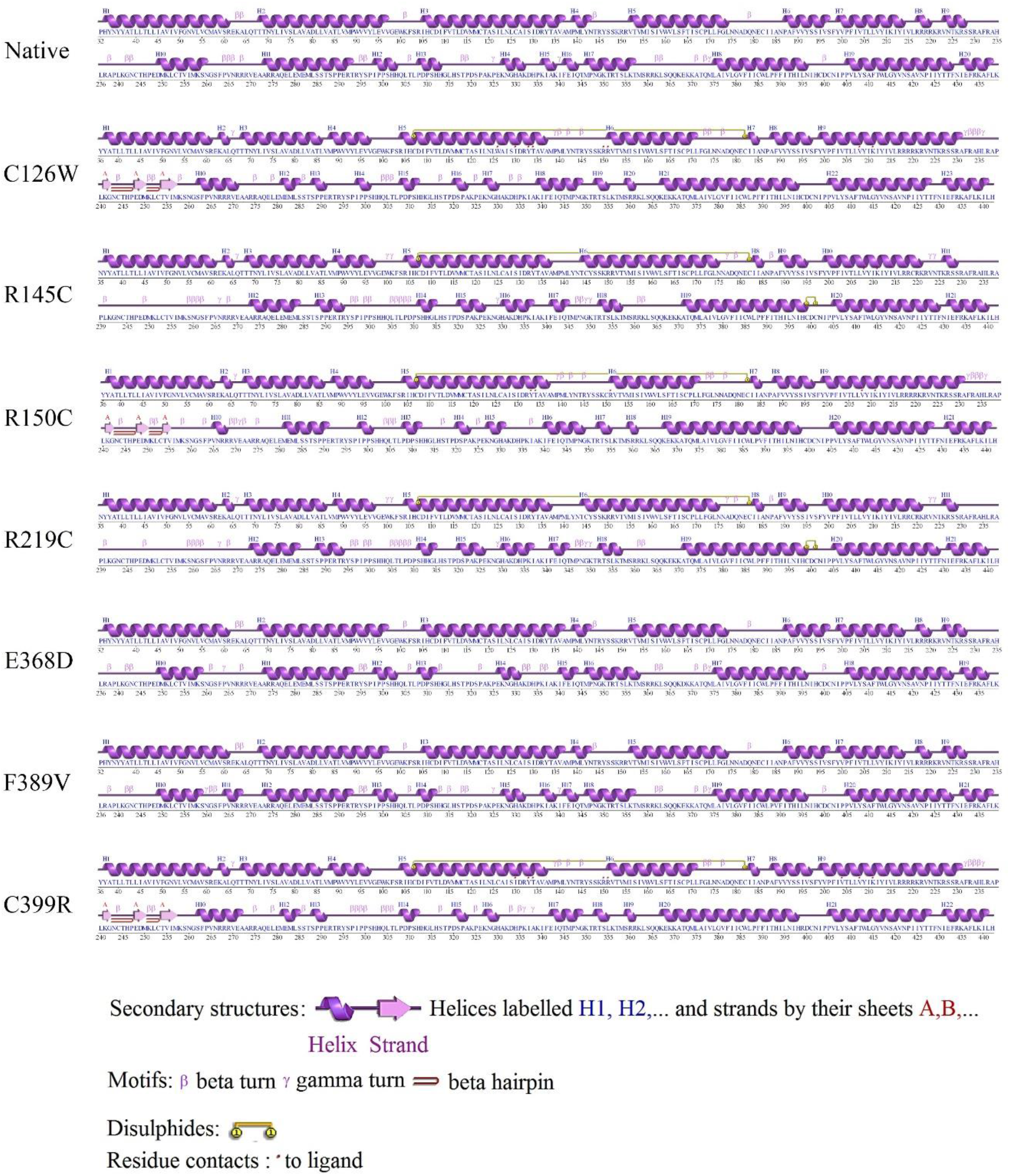
Secondary structure analysis of native DRD2 and the mutants revealed by PDBsum. It shows the modifications in terms of alpha helices, beta strands as well as various motifs that had occurred as a result of nsSNPs.

### Interatomic interactions prediction

Seven proteins selected from upstream analyses were analyzed using the DynaMut server (Fig. 5, Supplementary Table S6). The ΔΔG and Δ vibrational entropy energy predictions by ENCoM between the wild-type and mutant were displayed by DynaMut server. Here, the ΔΔG EnCoM value decreased in all mutants except for C126W and R150C in comparison with the wild-type. On the other hand, the ΔΔS ENCoM value in comparison with the wild-type protein increased in all candidates except for C126W and R150C. Notably, DynaMut ΔΔG predicted a total of 4 mutants (R145C, R219C, E368D, F389V) to be destabilizing all of which were detected by ENCoM to have increased molecule flexibility and decreased stability.

**Figure 5.**
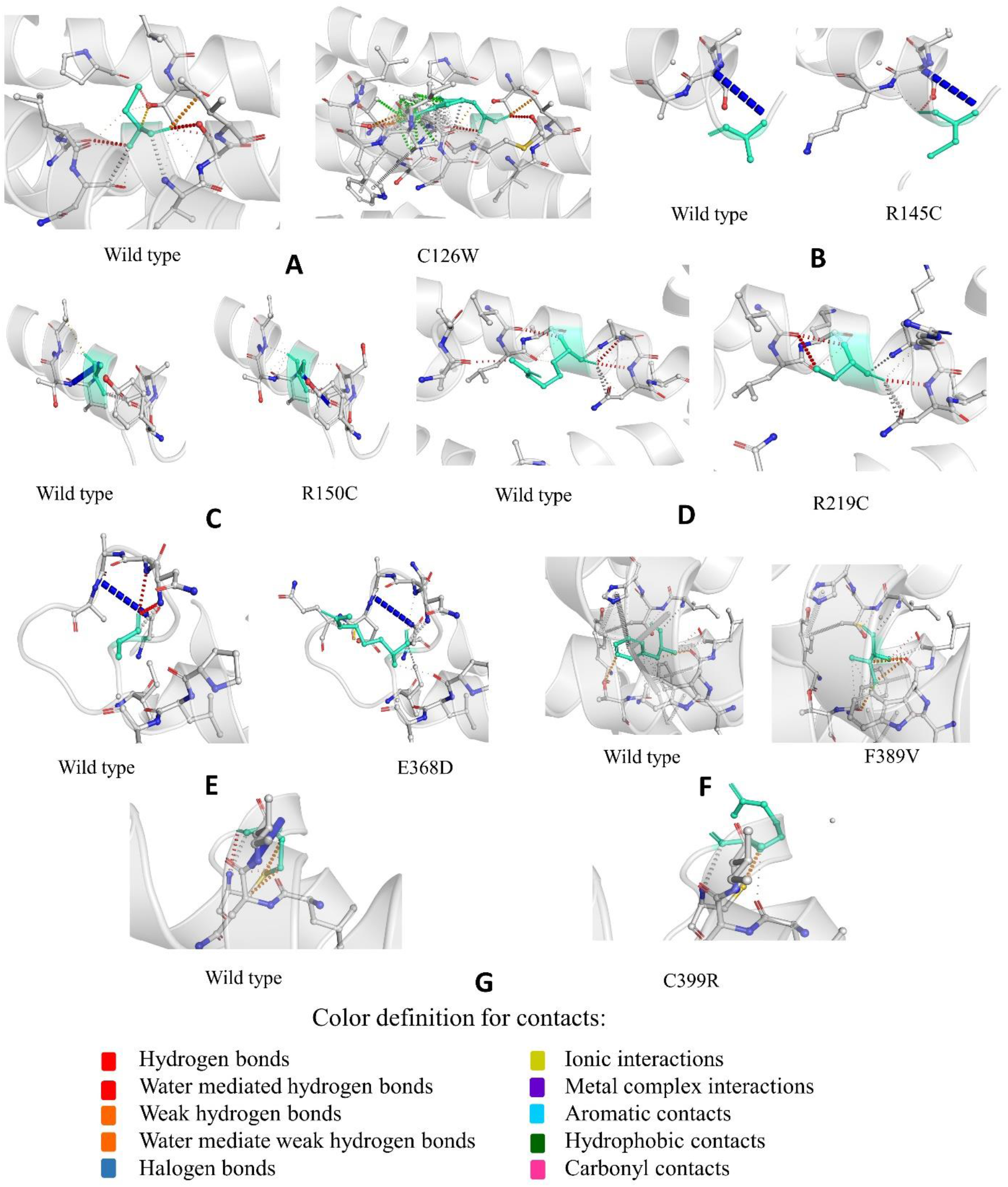
Dynamut prediction of inter-atomic interactions of the native DRD2 vs the mutants. Native and mutant residues are colored in light-green and showed as sticks along with the surrounding residues involved in interaction. Dot points with several colors represent the interactions such as hydrogen bonds, ionic interactions etc.

### Prediction of post-translational modification (PTM) sites

Post-translational modifications(PTMs) on protein can regulate diverse cellular processes including genotoxic stress response, innate immunity, and other cell signaling pathways ^62–64^. To reflect on the effect of the eight high-risk nsSNPs on PTMs, we used fourteen different *in silico* tools predicting probable PTM sites in the protein (Supplementary Table S7).

Upto 30% of mammalian proteins are estimated to be phosphorylated. Phosphorylation, ubiquitination, SUMOylation act as regulatory elements for diverse cellular functions such as protein conformational change, cellular division, differentiation, DNA repair growth, metabolism, immunity, learning and memory ^65–69^. For prediction of putative phosphorylation sites in DRD2 protein at serine, threonine and tyrosine residues, two servers GPS 5.0 and NetPhos 3.1 were used. Ubiquitination was predicted by UbPred and BDM-PUB whereas SUMOylation was predicted by GPS-SUMO 1.0.1 and SUMOplot servers. Other PTMs such as methylation, acetylation, and glycosylation have a regulatory role in many cellular pathways including metabolism, and homeostasis ^70^. Methylation sites were predicted by PSSMe, PMeS, and PLMLA tools.

Among all the residues identified by these fourteen different tools, only the residue R219 predicted by NetPhos 3.1 coincided with the filtered out high-risk SNPs.

### Molecular Docking analysis and visualization following binding site prediction

For the purpose of molecular docking analysis, at first, we used FTSite ligand binding site prediction tool. There were three sites found-two were in membrane embedded-extracellular part (binding site I and II) and one was in the cytosolic part (binding site III) of the protein (Fig. 6a). The reason for choosing Binding site II for molecular docking was that the residues in this site had been linked to dopamine and DRD2 antagonist (risperidone) binding by previous computational and crystal structure studies ^71,72^. From the 4 damaging nsSNPs filtered out from Dynamut, we chose only one mutation for the next steps. The reason for choosing F389V here was two-fold; one, while analysis by Project HOPE server it has been detected to situate within the agonist binding region and two, an earlier study ^72^ revealed this position to be involved in agonist binding.

**Figure 6.**
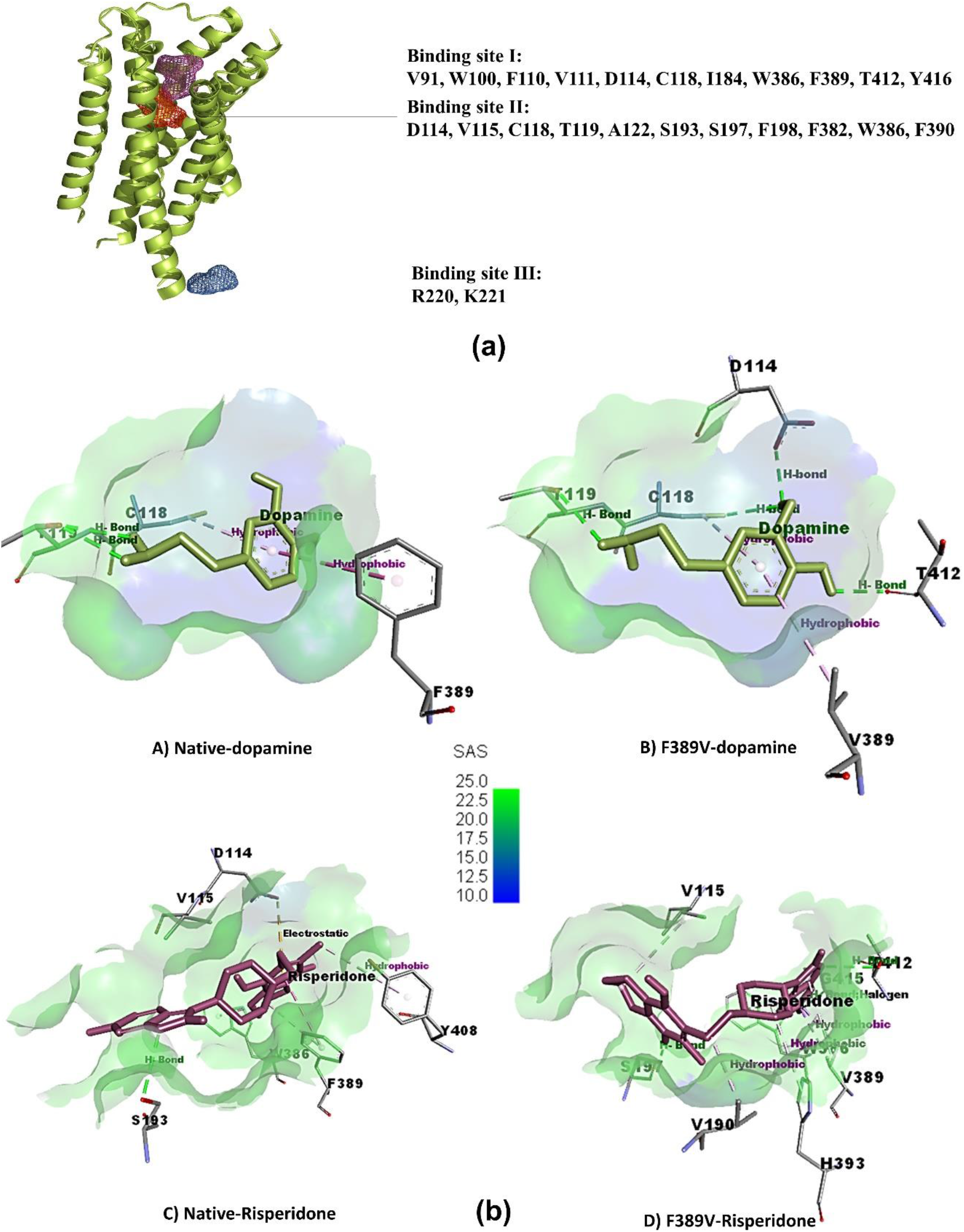
Binding site prediction and visualization of the molecular docking output. (a) FTSite predicted three ligand binding sites in DRD2. (b) Ligand non-bond binding interactions between the protein residues and the ligands visualized by Discovery Studio. Dopamine docked against (A) native DRD2 (B) F389V mutant; and risperidone docked against (C) native DRD2 and (D) F389V mutant are presented. The receptor surface is colored according to solvent accessibility surface (SAS) in an ascending degree from bule to green.

Following protein preparation by SwissPDB viewer and ligands by Avogadro software, a total of 4 molecular docking were carried out using AutoDock Vina in PyRx software (Supplementary Table S8). Residues in binding site II were specified for PyRx gridbox region. Binding affinity of natural agonist dopamine with wild-type (−5.8 kcal/mol) and mutant (−5.4 kcal/mol) shows that, mutation causes dopamine to bind with DRD2 less stably. Contrarily, the exact opposite scenario is observed in case of binding with the risperidone, a D2 receptor antagonist used as antipsychotic drug (wild-type −8.5kcal/mol and mutant −8.9kcal/mol) indicating an emergence of variability in drug response in case of a mutation. When visualized in Discovery Studio, docking interactions showed significant differences in mutant one from wild-type (Fig. 6b). When binding with dopamine, in comparison to native one, F389V mutated protein lost one hydrogen bond in T119 position and gained 3 hydrogen bonds in D114, C118 and T412 positions. Interaction with risperidone is somewhat complex for both wild-type and mutated proteins. While interacting, mutant lost electrostatic bond in D114 residue, one hydrogen bond in S193 and one hydrophobic bond in Y408. On the other hand, mutant protein gained two hydrogen bonds (S197, T412) and three hydrophobic bonds V190, W386 and H393) comparing to native structure. Apart from attaining one hydrogen bond, G415 position in mutant protein interacted with risperidone via a halogen bond too.

### Molecular Dynamics (MD) simulation

Since in physiological condition protein is dynamic in nature, molecular docking cannot give a holistic idea of protein behavior. Therefore, 200ns molecular dynamics simulation was carried out using the GROMACS software. (Fig. 7) The root-mean-square deviation (RMSD) analysis pointed that the F389V-dopamine complex underwent larger structural rearrangement compared to its native counterpart while the trend was opposite in case of F389V-risperidone complex. The dopamine and risperidone complex with the wild-type DRD2 were also found to attain stability faster than those with the F389V mutant. (Fig. 7a) The root-mean-square fluctuation (RMSF) values of each residue were analyzed since it reveals the rigid and flexible regions of a protein chain. The wild-type DRD2-dopamine complex had a total of 3 significant fluctuations (regions 210-220, 260-270, 290-300) while the F389V-dopamine complex exhibited total 2 but much larger fluctuations (regions 210-220, 300-325). In contrast, wild-type DRD2-risperidone complex displayed more frequent and larger peaks (regions 210-220, 230-240, 260-270, 300-310) compared to the mutant F389V-risperidone complex (regions 200-210, 300-310, and 320-330). (Fig. 7b) The radius of gyration (Rg) measurement was carried out to determine the degree of compactness of the protein. Rg of the wild-type DRD2-dopamine complex descended at a steady rate since the beginning of the simulation while the same for the F389V-dopamine complex elevated for 50 ns and then started to decrease. (Fig. 7c) For the wild-type DRD2-dopamine complex, the solvent accessible surface area (SASA) was at its peak at the beginning and had continued to decrease ever since. Meanwhile, the SASA for the F389V-dopamine complex increased for 50 ns and then started to drop. However, the F389V-risperidone complex displayed a stable decline in solvent accessibility while the wild-type DRD2-risperidone complex went through large fluctuations throughout the simulation. (Fig. 7d)

**Figure 7.**
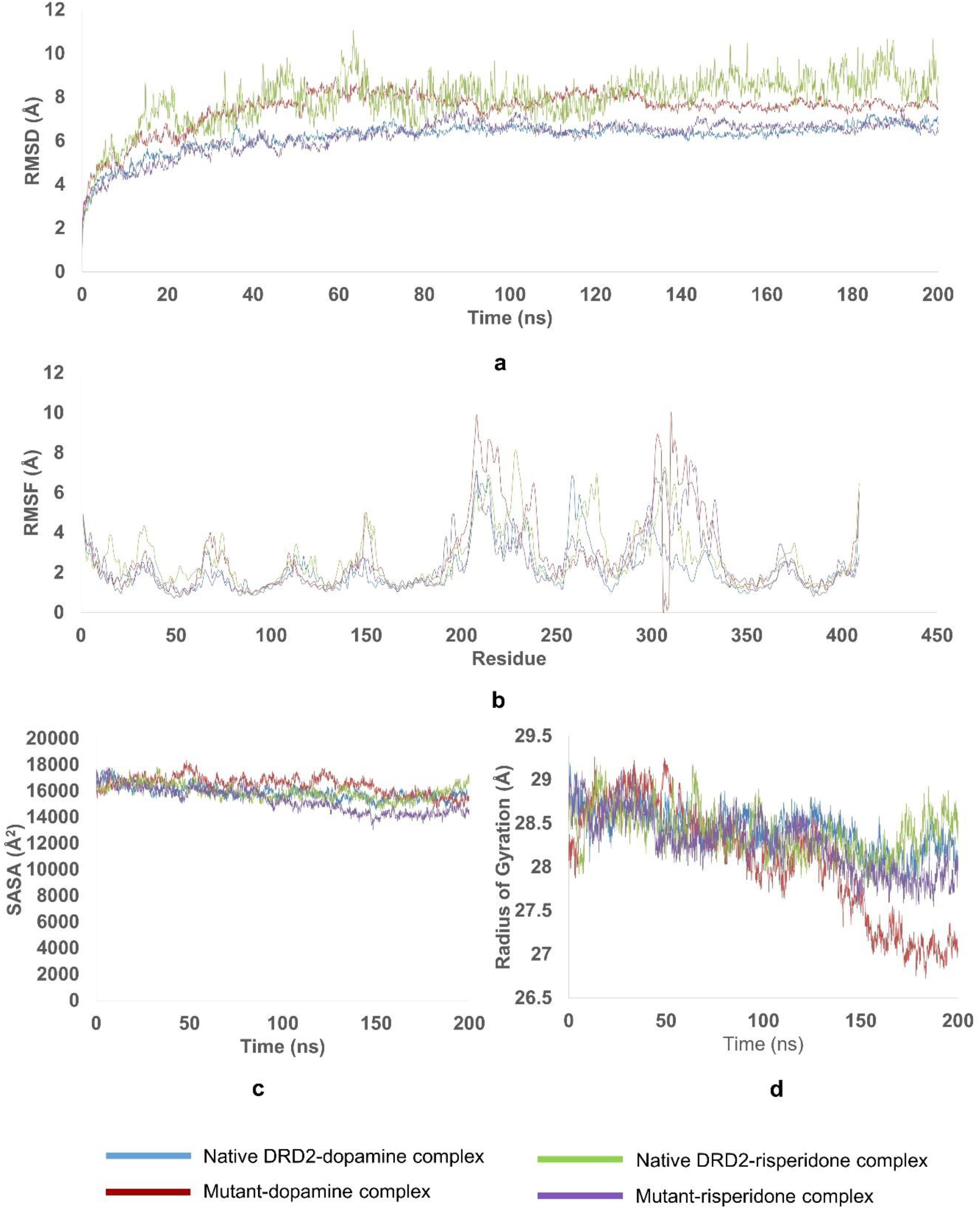
Results of 200 ns Molecular Dynamics Simulation by GROMACS software. Of the wild-type and F389V complexes with dopamine and risperidone: (a) RMSD values (b) RMSF values (c) SASA calculations and (d) Radius of gyration.

## Discussion

The present study aimed to investigate the effect of nsSNPs of *Drd2* on functional impairment and heterogeneity of psychotropic drug response. We pinpointed one mutation located within the agonist binding region-F389V, which in comparison with native protein, showed weaker binding affinity towards dopamine but a stronger affinity for the DRD2-antagonist risperidone.

Inter-individual variation in genetic components has the potential to influence not only disease susceptibility but also therapeutic response and drug-induced adverse effects. Studies on family, twin, pedigree, and epidemiological aspects support the idea of genetic association with psychiatric disorders, but they fail to provide informative data regarding psychotropic drug response. Since pharmacogenetic approaches consider individual subject’s genotypes, it provides an opportunity to identify biological predictors of psychotropic drug response by focusing exactly on the molecular substrates of drug activity ^73^.

Although there has been a number of studies on the associations between D2 receptor polymorphisms and neuropsychiatric disorders, the pharmacological aspects of the receptor polymorphisms are comparatively less attended. Essentially all antipsychotics clinically used for schizophrenia and bipolar disorder having the ability to block D2 dopamine receptors makes it a prime target of pharmacotherapy. A study by Healy and McKeon on dopamine D2-receptors and selective serotonin reuptake inhibitors (SSRIs) suggests that genetic factors affect response time to antidepressants and thereby can contribute to the variability in response time among patients ^74^. Sufficiency in antipsychotic binding affinity to D2 receptor for treatment efficacy is evidenced by correlation and functional imaging studies. A growing body of data suggests that D2 receptor polymorphisms influence response to clozapine, haloperidol, risperidone, and other neuroleptics. *Taq* A1 allele has been associated with early response to treatment with the antipsychotic nemonapride in schizophrenia. The same allele is also reported to function as a contributing factor for hyper-prolactinemia-related side effects in female patients by increasing prolactin levels ^75,76^.

Primarily, we sorted nsSNPs out based on their probable impact on functionality and structure. Different bioinformatics tools have their own threshold cut-off value for SNP categorization (as damaging and benign), which may sometimes result in false prediction for an SNP having a prediction score around the threshold cut-off value. We overcame this bias using a total of 15 tools based on sequence homology and structural homology approach of SNP prediction. Here is to mention that we have used the sequence of D2L isoform for the analyses. Because, unlike the D2S isoform, D2L is predominantly postsynaptic and antipsychotics (such as-risperidone, haloperidol) are evidenced to exert their effects by blockade of postsynaptic D2 receptors ^77,78^.

The number of SNP has been narrowed down from 260 to 9 after 2 steps analysis. All selected 9 nsSNPs were in the seven-transmembrane domain of the protein. All of the positions except for P404 are evolutionarily conserved indicating their role in protein structural stability. Following homology modeling, model verification, secondary structure analyses, and inter-atomic interaction prediction the list had been narrowed down to only 4 mutations – R145C, R219C, E368D and F389V.

Though the consequences of PTMs in DRD2 have not been reported yet, the regulatory role of PTMs in general are well-established. The arginine residue in 219 position is predicted by NetPhos 3.1 to be a potential phosphorylation site. As this residue is located within the region of interaction with the secondary messenger Neurabin 2 according to Project HOPE server and one molecular dynamics study ^79^, nsSNP R219C might impair the corresponding signaling cascade pathway.

Interestingly, one of these four mutations (F389V) was found to be located within the agonist binding region according to HOPE server. This property made it an ideal candidate to perform molecular docking with ligands such as agonists and antagonists. Although the charge and hydrophobicity of the F389V mutated protein remained unchanged according to the prediction by the HOPE server, the size was reported to decrease. Moreover, ConSurf detected this position of the native protein to be buried and involved in protein structure building while NetSurfP predicted conformational change of the F389V from buried to exposed. Besides, secondary structure attained by PDBsum showed that the mutant contained similar pattern of structure to native form upto N259 position but afterwards it had its β and γ turns more tightly packed. Dynamut server found it to be structurally destabilizing and more flexible.

PyRx was used for subsequent analysis by molecular docking followed by binding site prediction by FTSite. Among the 3 binding sites predicted by FTSite, two membrane-bound sites seem potential agonist-binding sites. Between these two, binding site II had more residues in common with the prediction from other *in silico* and *in vitro* studies on DRD2 agonist binding. Therefore, residues in this site were selected while performing molecular docking ^71,72^. We performed molecular docking with two ligands; one, DRD2 agonist-dopamine and two, a DRD2 antagonist-risperidone. Risperidone is a second-generation antipsychotic which has approximately 50-fold greater affinity for D2 receptor than that of clozapine but in low dose it is exclusive of side effects such as extrapyramidal symptoms (EPS) ^80,81^.

Insights from molecular docking and MD simulation inferred that the mutation F389V significantly affects the stability of the DRD2 protein. Binding affinity calculation of PyRx analysis showed that the F389V mutation decreased affinity of the receptor for dopamine but it made the receptor comparatively more responsive to risperidone. Non-bonding interactions analyses by Discovery Studio following molecular docking showed in the mutant F389V that interaction remained unchanged at the 389^th^ residue position, but changes were apparent in other residues involved in agonist binding. Although while binding with risperidone, the native protein lost electrostatic bond at position 114, a net gain of 1 hydrogen, 2 hydrophobic and 1 halogen bond made the bonding stronger in mutant protein. In case of dopamine binding, mutation caused the hydrogen bond to shift from 119 to other positions imposing impact on binding affinity and solvent accessibility. Findings from the molecular dynamics simulation reinforces that of molecular docking. The RMSD, RMSF and Rg values reveal that the mutation destabilized protein compactness and increased regional flexibility while in a complex with dopamine. However, the case was found to be exactly opposite while binding with risperidone. SASA calculations on dopamine complex suggested that the stability of the hydrophobic core of the protein was impaired in the F389V-dopamine complex. Contrarily, the mutation enhanced stability of the hydrophobic core while binding with risperidone. In brief, when the protein binds to its natural ligand, dopamine, the mutation reduces its stability. On the other hand, the mutation plays a positive role in stability while binding to the drug risperidone.

Our study has gone some way towards enhancing our understanding about polymorphism of the dopamine D2 receptor. Further experimental investigations are necessitated to elucidate the molecular and cellular basis of the observations. Apart from genetic variation, future studies should consider to include other factors potentially influencing interaction with D2 receptor, such as environment, age, body mass index of subjects etc. In case of sexually dimorphic diseases such as depression and ADHD, sex of subject is also a considerable factor. Furthermore, our findings give rise to the possibility of existence of other variants with similar effects. Broader scale of human *in vivo* molecular imaging studies is required to identify the most influential variants.

## Conclusion

Our study identified a potential high-risk variant of *Drd2* gene paving the way for extensive future studies on the nsSNPs of the gene. Further insights on drug targeting and biomarkers can be attained with the aid of *in vivo* models, GWAS and clinical studies.

## Materials and Method

### SNP dataset collection

The SNP data for human *Drd2* and the protein sequence were collected from NCBI dbSNP and UniProt database (UniProtKB ID P14416) (https://www.uniprot.org/) respectively ^82,83^.

### Functional impact prediction

Eight tools were employed for predicting the functional consequence of nsSNPs on DRD2 namely SIFT, PROVEAN, PolyPhen-2, SNAP 2, PANTHER-PSEP, P-Mut, SNPs & GO, PhD-SNP.

Depending on the physical properties and sequence homology of residues, SIFT (https://sift.bii.a-star.edu.sg/) predicts the effect of an amino acid substitution ^84^. Tolerance index score of SIFT is ≤0.05. PROVEAN (http://provean.jcvi.org/index.php) determines functionally important variants using a versatile alignment-based score ^85^. PROVEAN considers an nsSNP as deleterious when its score ≤–2.5. Both SIFT and PROVEAN take genomic co-ordinates of an SNP as the input. FASTA sequence of a protein and the location of SNP have to be provided as the input for the next 4 tools namely, PolyPhen-2, SNPs&GO, PANTHER and, Pmut. Utilizing physical and comparative considerations, PolyPhen-2 (http://genetics.bwh.harvard.edu/pph2/) predicts potential functional impact of an amino acid substitution on a protein ^86^. It classifies SNPs depending on probabilistic scores: possibly damaging (score >0.15), probably damaging (score >0.85), and benign (remaining). PANTHER (http://www.pantherdb.org/tools/csnpScoreForm.jsp) calculates the duration of evolutionary preservation of a residue, while longer duration indicates greater likelihood of harmful effect caused by mutation ^87^.

PMut (http://mmb.irbbarcelona.org/PMut/) is a web-based tool of which prediction score >0.5 indicates nsSNPs to have a damaging impact on protein function ^88^. The SNP & Gene Ontology (SNPs&GO) tool (https://snps.biofold.org/snps-and-go/snps-and-go.html) differentiates the disease associated variants from the benign ones based on protein sequence, gene ontology annotation and evolutionary patterns ^89^. As a neural network-based classifier, SNAP2 (https://rostlab.org/services/snap2web/) differentiates between impactful and neutral nsSNPs. The prediction score ranges from −100 to +100 meaning strong neutral to strong effect prediction respectively ^90^. PhD-SNP (https://snps.biofold.org/phd-snp/phd-snp.html) is a SVM-based classifier which uses either sequence based or sequence plus profile based algorithms ^91^.

### Structural impact prediction

Seven different structural impact prediction tools (MutPred, MUpro, NetSurf P, DUET, mCSM, SDM and HOPE) were used to find out structural impact.

MutPred (http://mutpred.mutdb.org/) provides a probabilistic idea about the impact of amino acid substitutions via a machine learning-based method ^92^. Using Support Vector Machines and Neural Networks, MUpro (https://www.ics.uci.edu/~baldig/mutation.html) predicts protein stability changes for point mutations with about 84% accuracy ^93^. Both MutPred and MUpro takes protein FASTA sequence for input. NetSurfP https://services.healthtech.dtu.dk/service.php?NetSurfP-2.0) is a sequence-based local structural feature prediction server giving idea about solvent accessibility for input residues ^94^.

DUET, mCSM and SDM (http://biosig.unimelb.edu.au/duet/) are closely intertwined tools and their input can be provided together in the form of the native protein sequence while separately mentioning the relevant SNP position. 14–16 Project HOPE server (http://www.cmbi.ru.nl/hope/input/) anticipates the structural impacts of nsSNPs by integrating information from a wide array of sources including tertiary structure, sequence annotations, homology models from the Distributed Annotation System (DAS) servers and UniProt database etc. 17 It takes the native protein sequence as the input and asks for the specific site and type of mutation to be analyzed before proceeding to analysis.

### Identification of domain

Utilized databases for identification of conserved domain and positions of nsSNPs in domain are – InterPro (http://www.ebi.ac.uk/interpro/), Pfam (http://pfam.xfam.org/), PROSITE (https://prosite.expasy.org/prosite.html) and CDD (https://www.ncbi.nlm.nih.gov/cdd/) ^95–98^. InterPro is a powerful tool which combines several databases while Pfam data is based on the UniProt Reference Proteomes. PROSITE works by grouping of similarities in protein and CDD uses position-specific score matrices ([PSSMs](https://www.ncbi.nlm.nih.gov/Structure/cdd/cdd_help.shtml#CD_PSSM)) for domain prediction. Protein FASTA sequence was given as input for all four cases.

### Evolutionary conservation analysis

To unravel the evolutionary history of the human DRD2 protein, a phylogenetic tree was constructed in MEGA X tool using the 10 closest matches to the human DRD2 protein as determined by a BLASTp search (https://blast.ncbi.nlm.nih.gov/Blast.cgi?PAGE=Proteins) ^99^. The maximum likelihood algorithm and bootstrap value of 1000 was used to build the tree. It was then visualized with the Iroki webserver (https://www.iroki.net/) ^100^.

ConSurf (https://consurf.tau.ac.il/) provides conservation analysis of individual amino acids in the protein based on the phylogenetic relationships between homologous sequences ^101^. Upon providing the PDB structure of query protein, the server output was found to be in 3 classes: score 1-4 indicates variable, 5-6 is considered intermediate and 7-9 means conserved. HOPE server was also employed for generating for each nsSNP that had been filtered through the previous steps.

### Homology model verification and secondary structure analysis

Upon generation 3D structure of native and mutant proteins via SWISS-MODEL (https://swissmodel.expasy.org/), structure quality was exposed to verification by PROCHECK and ERRAT. ERRAT provided a quality score while Ramachandran plots were generated from PROCHECK. In general, a protein structure is considered good if more than 90% residues are in the favored region of Ramachandran plot. PDBsum (http://www.ebi.ac.uk/thornton-srv/databases/cgi-bin/pdbsum/GetPage.pl?pdbcode=index.html) provides features of secondary structure such as alpha helices, beta strands, beta hairpins etc of proteins taking amino acid sequences as input ^102^.

### Interatomic interactions prediction

DynaMut server was used to predict the changes in interatomic interactions interaction upon point mutation (http://biosig.unimelb.edu.au/dynamut/) ^103^. Wild-type structure in PDB format and mutation list were provided as input.

### Predicting post-translational modification (PTM) sites

Different post-translational modification (PTM) sites, including Phosphorylation, Ubiquitination, SUMOylation, Methylation, Acetylation and Glycosylation were predicted by fourteen different online servers. Both GPS 5.0 (http://gps.biocuckoo.cn/) and NetPhos 3.1 (http://www.cbs.dtu.dk/services/NetPhos/) predicted phosphorylation of serine, threonine and tyrosine residues ^104,105^. While GPS 5.0 works using position weight determination (PWD) and scoring matrix optimization (SMO) methods, NetPhos 3.1 uses ensembles of neural networks to perform the prediction. For GPS 5.0, threshold was kept medium whereas it was 0.5 for Netphos 3.1. Of the two servers used for ubiquitination prediction, UbPred is a forest-based predictor and BDM-PUB utilizes Bayesian Discriminant method. In UbPred (http://www.ubpred.org/), the lysine amino acid having a score of 0.62–69, 0.69–0.84, and 0.84–1 were assigned as low, medium, and high confidence ubiquitination sites respectively ^106^. In case of BDM-PUB (http://bdmpub.biocuckoo.org/index.php), threshold was 0.3 and balanced cut-off option was selected. Both the servers GPS-SUMO and SUMOplot utilized manually curated dataset; GPS-SUMO 1.0.1 employed the method Particle Swarm Optimization (PSO) but SUMOplot used MotifX software. GPS-SUMO 1.0.1 (http://sumosp.biocuckoo.org/) threshold were kept medium and cut-of value range was 16-59.29 and only high probability motifs with a score >0.5 (0.79, 0.67) were considered SUMOylated in SUMOplot (https://www.abcepta.com/sumoplot) ^107^. NetAcet 1.0 (http://www.cbs.dtu.dk/services/NetAcet/) predicted acetylation in Alanine, Glycine, Serine and Threonine residues ^108^. For Netglycate 1.0 (http://www.cbs.dtu.dk/services/NetGlycate/), scores greater than 0 indicate glycation sites while the score range is–1 to 1 ^109^. Threshold for PSSMe (http://bioinfo.ncu.edu.cn/PSSMe.aspx), PMeS (http://bioinfo.ncu.edu.cn/inquiries_PMeS.aspx), PLMLA (http://bioinfo.ncu.edu.cn/inquiries_PLMLA.aspx), NetNGlyc (http://www.cbs.dtu.dk/services/NetNGlyc/) and NetOGlycate 4.0 (http://www.cbs.dtu.dk/services/NetOGlyc/) is 0.5. respectively ^110–113^.

For all the tools the FASTA protein sequence of DRD2 was submitted except for PSSMe which needed UniProt id also.

### Molecular Docking analysis and visualization following binding site prediction

FTSite (https://ftsite.bu.edu/) recognizes ligand binding sites on proteins taking PDB structure as input ^114^.

The proteins and ligands were energy minimized using SwissPDB viewer and Avogadro respectively. ^115,116^ Then the AutoDock Vina embedded within the PyRx molecular docking software was used to dock the respective proteins and the ligands. ^117^ All the parameters were set as default and the grid box covered the agonist binding site as predicted by FTSite. The graphical output of the docked complexes were subjected to in-depth analysis in Discovery Studio for revealing the crucial information about interactions between the proteins and the ligands ^118^.

### Molecular Dynamics (MD) simulation

MD simulation for 200 ns was carried out using GROningen MAchine for Chemical Simulations aka GROMACS (version 5.1.1) ^119^. The GROMOS96 43a1 force-field was applied. The physiological condition of the system was defined as (300 K, pH 7.4, 0.9% NaCl). The structures were solvated in a dodecahedral box of the SPC (simple point charge) water model with its edges at 1 nm distance from the protein surface. The overall charge of the system was neutralized through the addition of 24 Cl^-^ions using the genion module. Energy minimization of the neutralized system was carried out using the steepest descent minimization algorithm. The ligand was restrained before carrying out the isothermal-isochoric (NVT) equilibration of the system for 100 ps with short-range electrostatic cutoff value of 1.2 nm. Following NVT, Isobaric (NPT) equilibration of the system was carried out for 100 ps with short-range van der Waals cutoff fixed at 1.2 nm. Finally, a 200 ns molecular dynamic simulation was run using periodic boundary conditions. The energy of the system was saved every 100 ps. For calculating the long range electrostatic potential, the Particle Mesh Ewald (PME) method was applied. Short-range van der Waals cutoff was kept at 1.2 nm. Modified Berendsen thermostat was used to control simulation temperature while the pressure was kept constant using the Parrinello-Rahman algorithm. The simulation time step was selected as 2.0 fs while the snapshot interval was set to 100 ps for analyzing the trajectory data. Upon successful completion of the simulation, all of the trajectories were concatenated to calculate and plot RMSD, RMSF, Rg and SASA data. The RMSD, RMSF, Rg and SASA calculations were carried out using the rms, rmsf, gyrate, and sasa modules respectively.

## Author contributions

S.S.L. and I.A. conceived the idea, designed the study, preformed analyses, wrote and critically revised the manuscript.

## Supplementary information

**Supplementary Table S1.** Distribution of STK11 intron, missense, coding synonymous, non-coding transcript variant and intron, 3’UTR, and 5’UTR SNPs.

| SNP | Amount |
| --- | --- |
| Intron | 15370 (93.40%) |
| Missense | 238 (1.45%) |
| Synonymous | 173 (1.05%) |
| Non-coding | 139 (0.84%) |
| In-frame deletion | 2 (0.01%) |
| Total | 16456 |

**Supplementary Table S2.**
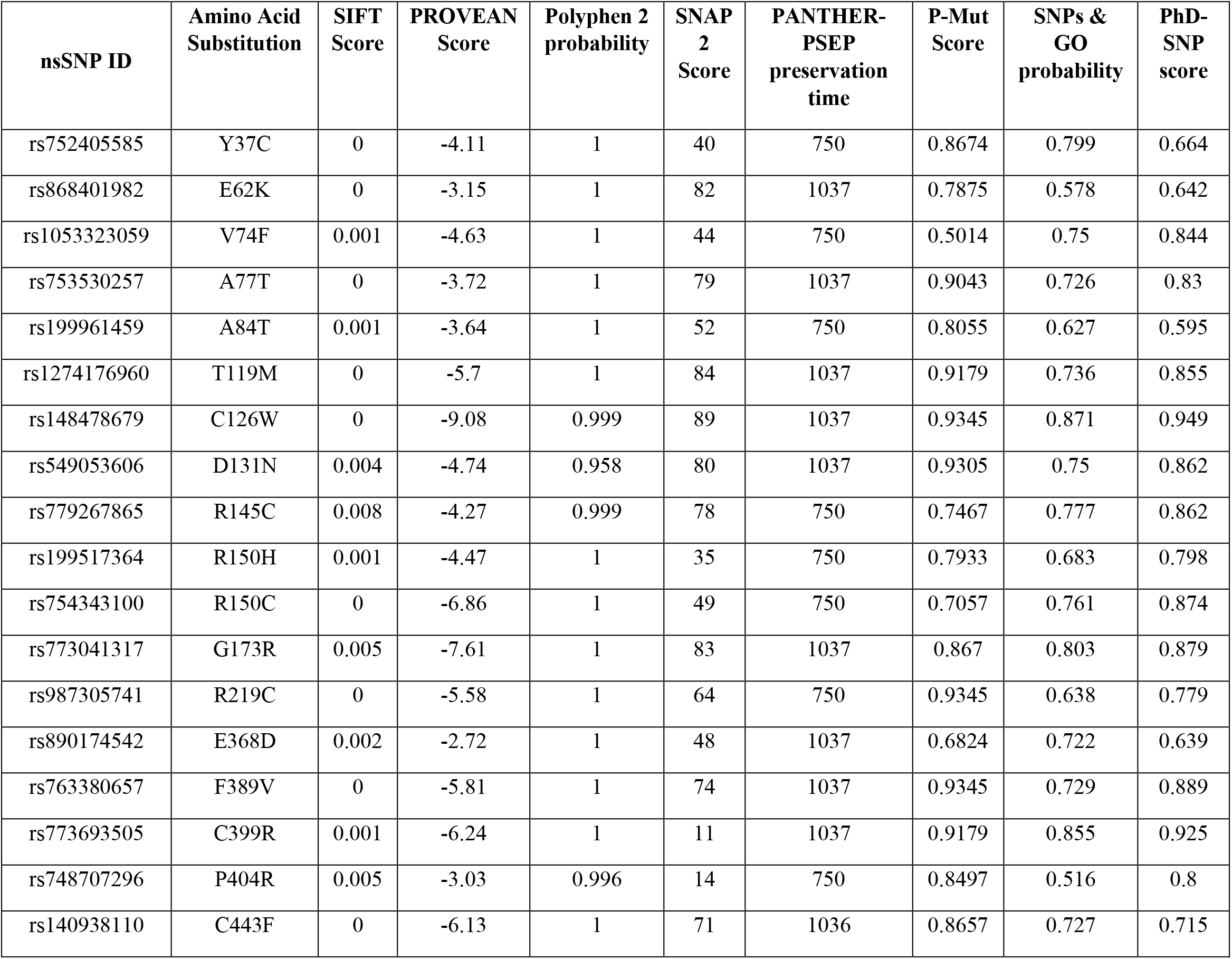
Unanimous deleterious or damaging missense SNPs in the DRD2 protein predicted by 8 different tools.

| nsSNP ID | Amino Acid Substitution | SIFT Score | PROVEAN Score | Polyphen 2 probability | SNAP 2 Score | PANTHER-PSEP preservation time | P-Mut Score | SNPs & GO probability | PhD-SNP score |
| --- | --- | --- | --- | --- | --- | --- | --- | --- | --- |
| rs752405585 | Y37C | 0 | -4.11 | 1 | 40 | 750 | 0.8674 | 0.799 | 0.664 |
| rs868401982 | E62K | 0 | -3.15 | 1 | 82 | 1037 | 0.7875 | 0.578 | 0.642 |
| rs1053323059 | V74F | 0.001 | -4.63 | 1 | 44 | 750 | 0.5014 | 0.75 | 0.844 |
| rs753530257 | A77T | 0 | -3.72 | 1 | 79 | 1037 | 0.9043 | 0.726 | 0.83 |
| rs199961459 | A84T | 0.001 | -3.64 | 1 | 52 | 750 | 0.8055 | 0.627 | 0.595 |
| rs1274176960 | T119M | 0 | -5.7 | 1 | 84 | 1037 | 0.9179 | 0.736 | 0.855 |
| rs148478679 | C126W | 0 | -9.08 | 0.999 | 89 | 1037 | 0.9345 | 0.871 | 0.949 |
| rs549053606 | D131N | 0.004 | -4.74 | 0.958 | 80 | 1037 | 0.9305 | 0.75 | 0.862 |
| rs779267865 | R145C | 0.008 | -4.27 | 0.999 | 78 | 750 | 0.7467 | 0.777 | 0.862 |
| rs199517364 | R150H | 0.001 | -4.47 | 1 | 35 | 750 | 0.7933 | 0.683 | 0.798 |
| rs754343100 | R150C | 0 | -6.86 | 1 | 49 | 750 | 0.7057 | 0.761 | 0.874 |
| rs773041317 | G173R | 0.005 | -7.61 | 1 | 83 | 1037 | 0.867 | 0.803 | 0.879 |
| rs987305741 | R219C | 0 | -5.58 | 1 | 64 | 750 | 0.9345 | 0.638 | 0.779 |
| rs890174542 | E368D | 0.002 | -2.72 | 1 | 48 | 1037 | 0.6824 | 0.722 | 0.639 |
| rs763380657 | F389V | 0 | -5.81 | 1 | 74 | 1037 | 0.9345 | 0.729 | 0.889 |
| rs773693505 | C399R | 0.001 | -6.24 | 1 | 11 | 1037 | 0.9179 | 0.855 | 0.925 |
| rs748707296 | P404R | 0.005 | -3.03 | 0.996 | 14 | 750 | 0.8497 | 0.516 | 0.8 |
| rs140938110 | C443F | 0 | -6.13 | 1 | 71 | 1036 | 0.8657 | 0.727 | 0.715 |

**Supplementary Table S3.** Structural impact prediction of DRD2 high-risk pathogenic nsSNPs of DRD2 protein.

| Amino Acid Substitution |  | Y37C | E62K | V74F | A77T | A84T | T119M | C126W | D131N | R145C |
| --- | --- | --- | --- | --- | --- | --- | --- | --- | --- | --- |
| Mutpred prediction |  | Benign | Pathogenic | Pathogenic | Pathogenic | Pathogenic | Pathogenic | Pathogenic | Pathogenic | Pathogenic |
| Mupro stability prediction |  | Decrease | Decrease | Decrease | Decrease | Decrease | Increase | Decrease | Decrease | Decrease |
| NetSurfP prediction | native | Buried | Exposed | Buried | Buried | Exposed | Buried | Buried | Buried | Exposed |
|  | mutant | Buried | Exposed | Buried | Buried | Buried | Buried | Buried | Buried | Exposed |
| mCSM Prediction |  | Destabilizing | Destabilizing | Destabilizing | Destabilizing | Destabilizing | Destabilizing | Destabilizing | Destabilizing | Destabilizing |
| SDM prediction |  | Stabilizing | Stabilizing | Destabilizing | Destabilizing | Destabilizing | Stabilizing | Stabilizing | Destabilizing | Destabilizing |
| DUET prediction |  | Destabilizing | Stabilizing | Destabilizing | Destabilizing | Destabilizing | Stabilizing | Destabilizing | Destabilizing | Destabilizing |
| Change in size |  | Decrease | Increase | Increase | Increase | Increase | Increase | Increase | Unchanged | Decrease |
| Change of charge |  | Unchanged | Negative > Positive | Unchanged | Unchanged | Unchanged | Unchanged | Unchanged | Negative > Neutral | Positive > Neutral |
| Change in hydrophobicity |  | Increase | Unchanged | Unchanged | Loss | Loss | Increase | Unchanged | Unchanged | Increase |
| Other impacts |  | None | None | None | None | None | None | The residue being involved in a cysteine bridge, the mutation will have a severe effect on the 3D-structure. | None | The residue being involved in a cysteine bridge, the mutation will have a severe effect on the 3D-structure. |

**Supplementary Table S3.** Structural impact prediction of DRD2 high-risk pathogenic nsSNPs of DRD2 protein. (continued)
| Amino Acid Substitution |  | R150H | R150C | G173R | R219C | E368D | F389V | C399R | P404R | C443F |
| --- | --- | --- | --- | --- | --- | --- | --- | --- | --- | --- |
| Mutpred score |  | Pathogenic | Pathogenic | Pathogenic | Pathogenic | Pathogenic | Pathogenic | Pathogenic | Pathogenic | Pathogenic |
| Mupro stability prediction |  | Decrease | Decrease | Decrease | Decrease | Decrease | Decrease | Decrease | Decrease | Increase |
| NetSurfP prediction | native | Exposed | Exposed | Exposed | Exposed | Exposed | Buried | Exposed | Exposed | Exposed |
|  | mutant | Exposed | Exposed | Exposed | Buried | Exposed | Exposed | Exposed | Exposed | Exposed |
| mCSM Prediction |  | Destabilizing | Destabilizing | Destabilizing | Destabilizing | Destabilizing | Destabilizing | Stabilizing | Destabilizing | Destabilizing |
| SDM prediction |  | Destabilizing | Destabilizing | Destabilizing | Destabilizing | Destabilizing | Stabilizing | Stabilizing | Destabilizing | Stabilizing |
| DUET prediction |  | Destabilizing | Destabilizing | Destabilizing | Destabilizing | Destabilizing | Destabilizing | Stabilizing | Stabilizing | Destabilizing |
| Change in size |  | Decrease | Decrease | Increase | Decrease | Decrease | Decrease | Increase | Increase | Increase |
| Change of charge |  | Positive > Neutral | Positive > Neutral | Neutral > Positive | Positive > Neutral | Unchanged | Unchanged | Positive > Neutral | Neutral > Positive | Unchanged |
| Change in hydrophobicity |  |  | Increase | Increase | Increase | Unchanged | Unchanged | Decrease | Decrease | Unchanged |
| Other impacts |  | None | The residue being involved in a cysteine bridge, the mutation will have a severe effect on the 3D-structure. | Exclusively the native glycine forms an unusual torsion angle. The mutation will force the local backbone into an incorrect conformation and will disturb the local structure. | The mutation will disturb interaction with secondary messenger Neurabin 2. | The wild-type forms a H bond with Gln 368 and a salt bridge with Lys 369. The mutation will disturb interaction with secondary messenger Neurabin 2. | The mutation being located within the agonist binding region, the resulting empty space in the core can disturb the functionality of the protein | The residue being involved in a cysteine bridge, the mutation will have a severe effect on the 3D-structure. | The mutation can disturb the special backbone conformation induced by the native proline residue. | None |

**Supplementary Table S4.** Evolutionary conservation prediction of DRD2 protein using ConSurf and HOPE server

| **Residue and Position** | **Conservation score** | **Prediction** | **HOPE conservation prediction** |
| --- | --- | --- | --- |
| C126 | 9 | Highly conserved and buried (s) | Very conserved |
| R145 | 8 | Very conserved and exposed (f) | Very conserved |
| R150 | 9 | Highly conserved and exposed (f) | Very conserved |
| G173 | 9 | Highly conserved and buried (s) | Very conserved |
| R219 | 9 | Highly conserved and exposed (f) | Very conserved |
| E368 | 9 | Highly conserved and exposed (f) | Very conserved |
| F389 | 9 | Highly conserved and buried (s) | 100% conserved |
| C399 | 9 | Highly conserved and buried (s) | Very conserved |
| P404 | 7 | Conserved and exposed | Not very conserved |

**Supplementary Table S5.**
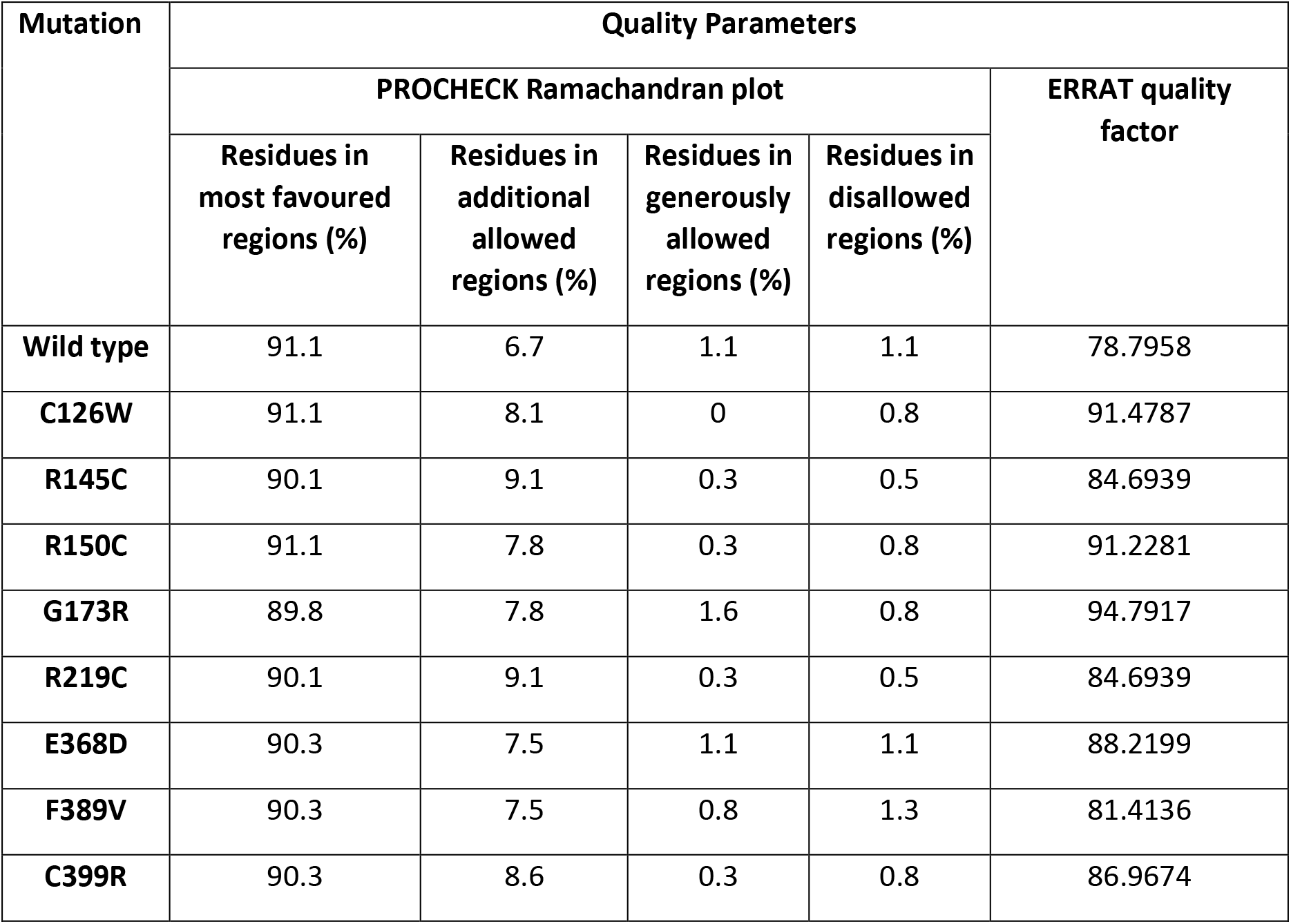
Protein model verification by PROCHECK and ERRAT.

| Mutation | Quality Parameters |  |  |  |  |
| --- | --- | --- | --- | --- | --- |
|  | PROCHECK Ramachandran plot |  |  |  | ERRAT quality factor |
|  | Residues in most favoured regions (%) | Residues in additional allowed regions (%) | Residues in generously allowed regions (%) | Residues in disallowed regions (%) |  |
| Wild type | 91.1 | 6.7 | 1.1 | 1.1 | 78.7958 |
| C126W | 91.1 | 8.1 | 0 | 0.8 | 91.4787 |
| R145C | 90.1 | 9.1 | 0.3 | 0.5 | 84.6939 |
| R150C | 91.1 | 7.8 | 0.3 | 0.8 | 91.2281 |
| G173R | 89.8 | 7.8 | 1.6 | 0.8 | 94.7917 |
| R219C | 90.1 | 9.1 | 0.3 | 0.5 | 84.6939 |
| E368D | 90.3 | 7.5 | 1.1 | 1.1 | 88.2199 |
| F389V | 90.3 | 7.5 | 0.8 | 1.3 | 81.4136 |
| C399R | 90.3 | 8.6 | 0.3 | 0.8 | 86.9674 |

**Supplementary Table S6.** Interatomic interaction prediction of native DRD2 and mutant proteins.

| Mutation | $\Delta\Delta G$<br>ENCoM | $\Delta\Delta S$<br>ENCoM | $\Delta\Delta G$<br>DynaMut |
| --- | --- | --- | --- |
| C126W | 0.55 | -0.688 | 1.229 |
| R145C | -0.174 | 0.218 | -0.512 |
| R150C | 0.403 | -0.504 | 0.412 |
| R219C | -0.34 | 0.425 | -0.621 |
| E368D | -0.267 | 0.334 | -0.838 |
| F389V | -0.655 | 0.819 | -0.206 |
| C399R | -0.501 | 0.626 | 0.054 |

**Supplementary Table S7.**
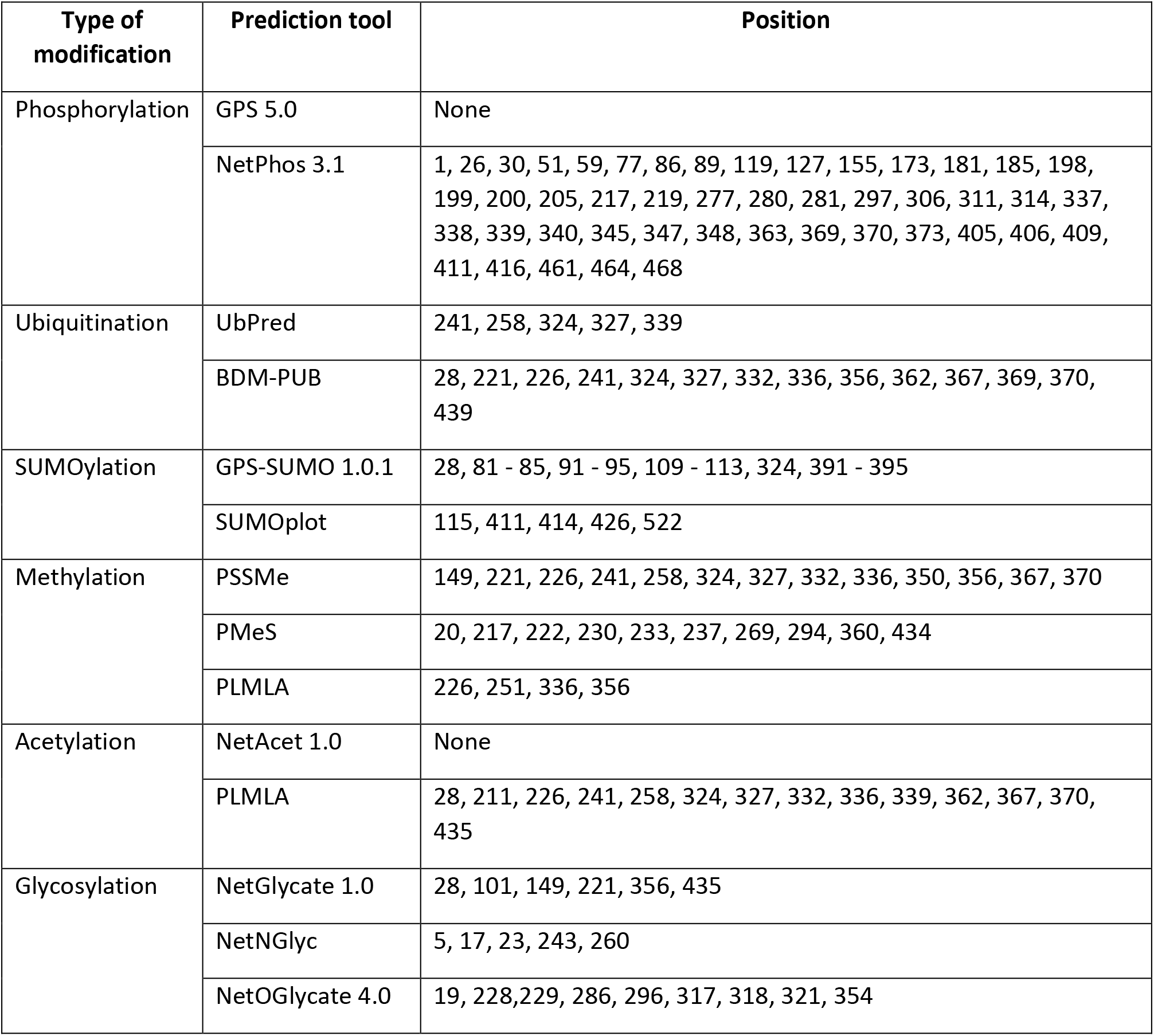
Post-translational modification site prediction in DRD2 protein.

**Supplementary Table S8.** Binding affinity (kcal/mol) prediction of ligands with native and mutant using PyRx.

| Ligand | Binding affinity (kcal/mol) |  |
| --- | --- | --- |
|  | Wild type | Mutant (F389V) |
| Dopamine | -5.8 | -5.4 |
| Risperidone | -8.5 | -8.9 |

**Supplementary Figure 1.**
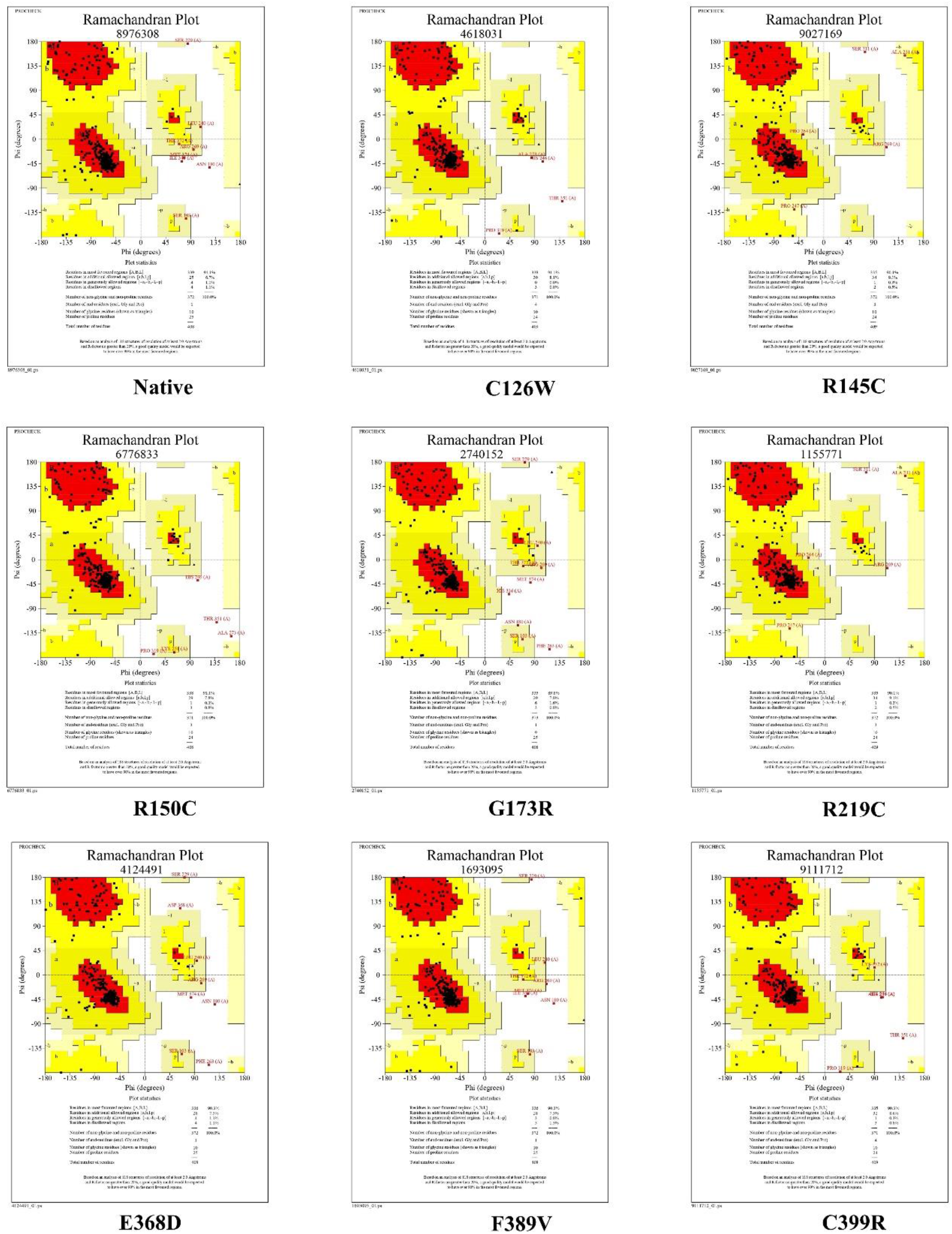
Ramachandran plots of native and mutants derived from PROCHECK.

